# Highly reduced genomes of protist endosymbionts show evolutionary convergence

**DOI:** 10.1101/719211

**Authors:** Emma E. George, Filip Husnik, Daria Tashyreva, Galina Prokopchuk, Aleš Horák, Waldan K. Kwong, Julius Lukeš, Patrick J. Keeling

**Affiliations:** University of British Columbia, Department of Botany, Vancouver, Canada; Institute of Parasitology, Biology Center, Czech Academy of Sciences, České Budějovice, Czech Republic; Faculty of Sciences, University of South Bohemia, České Budějovice, Czech Republic

**Keywords:** Endosymbiosis, genome reduction, convergent evolution, *Rickettsiaceae*, *Holosporaceae*, diplonemid, T6SS

## Abstract

Genome evolution in bacterial endosymbionts is notoriously extreme: the combined effects of strong genetic drift and unique selective pressures result in highly reduced genomes with distinctive adaptations to hosts [1–4]. These processes are mostly known from animal endosymbionts, where nutritional endosymbioses represent the best-studied systems. However, eukaryotic microbes, or protists, also harbor diverse bacterial endosymbionts, but their genome reduction and functional relationships with their more diverse hosts are largely unexplored [5–7]. We sequenced the genomes of four bacterial endosymbionts from three species of diplonemids, poorly-studied but abundant and diverse heterotrophic protists [8–10]. The endosymbionts come from two intracellular families from different orders, *Rickettsiaceae* and *Holosporaceae*, that have invaded diplonemids multiple times, and their genomes have converged on an extremely small size (605–632 kbp), similar gene content (e.g., metabolite transporters and secretion systems), and reduced metabolic potential (e.g., loss of energy metabolism). These characteristics are generally found in both families, but the diplonemid endosymbionts have evolved greater extremes in parallel. Their modified type VI secretion systems are likely involved in the manipulation of host metabolism (e.g., interactions with host mitochondria) or defense against bacterial infections, although their similar effector/immunity proteins may also allow for co-occurring *Holosporaceae* species in one diplonemid host. Finally, modified cellular machinery like ATP synthase without oxidative phosphorylation and reduced flagella present in both diplonemid endosymbionts and nutritional animal endosymbionts indicates that intracellular mechanisms have converged in bacterial endosymbionts with various functions and from different eukaryotic hosts across the tree of life.

## Results

### Diplonemid endosymbionts represent new, phylogenetically divergent species

Cultures of three diplonemid species where endosymbionts have been recorded and formally described [11,12] were screened to confirm the presence and identity of their full complement of endosymbionts by fluorescent in-situ hybridization (FISH) and sequencing of the bacterial 16S rRNA gene (see Star Methods). *Diplonema aggregatum* and *Diplonema japonicum* were confirmed to contain *Holosporaceae* species, *Cytomitobacter indipagum* and *Cytomitobacter primus*, respectively (bacterial taxa will be referred to without the *Candidatus* prefix) [11]. Interestingly, in *D. japonicum* a second and more abundant but previously undetected endosymbiont was found that was closely related to *Cytomitobacter* (Figure S1), but distinct enough to warrant a new genus (~87% 16S rRNA sequence identity to *Cytomitobacter*, well below the 94.5% gene sequence identity threshold for genera [13]) (Figure S2). We propose the new species, *Nesciobacter abundans* gen. nov., sp. nov. (from Latin *nescio* meaning ‘I do not know’ and *abundans* due to its higher abundance in the host), for this novel *Holosporaceae* endosymbiont (see full description below). Finally, the more distantly related diplonemid, *Namystynia karyoxenos*, was confirmed to contain the *Rickettsiaceae* species *Sneabacter namystus* [12] (Figure S1).

The relative abundance and distribution of *C. primus* and *N. abundans* were compared in *D. japonicum* using species-specific fluorescence *in situ* hybridization (FISH) probes. Both endosymbionts were present in all host cells we examined, where they were sporadically distributed beneath the host cell’s surface (Figure 1). The population size of both endosymbiont species and their ratio changed depending on the host life stage. The total abundance of bacterial endosymbionts was significantly higher and more variable in the larger trophic (feeding) hosts with 24 to 108 bacteria per host (mean ± SD [standard deviation of mean], 53.1 ± 17.8; n = 127) compared to swimming (starving) hosts, with 16 to 48 bacteria per host (32.7 ± 6.9; n = 120). Swimming cells maintained a relatively stable 7:3 ratio of *N. abundans* to *C. primus* cells across cells within and between replicate cultures (28.9 ± 6.5% *C. primus*; n = 120). In trophic hosts the ratio was more variable both among cells and across three replicates (23.9 ± 9.7% *C. primus*; n = 127), although *N. abundans* was more abundant in all observed host cells.

**Figure 1.**
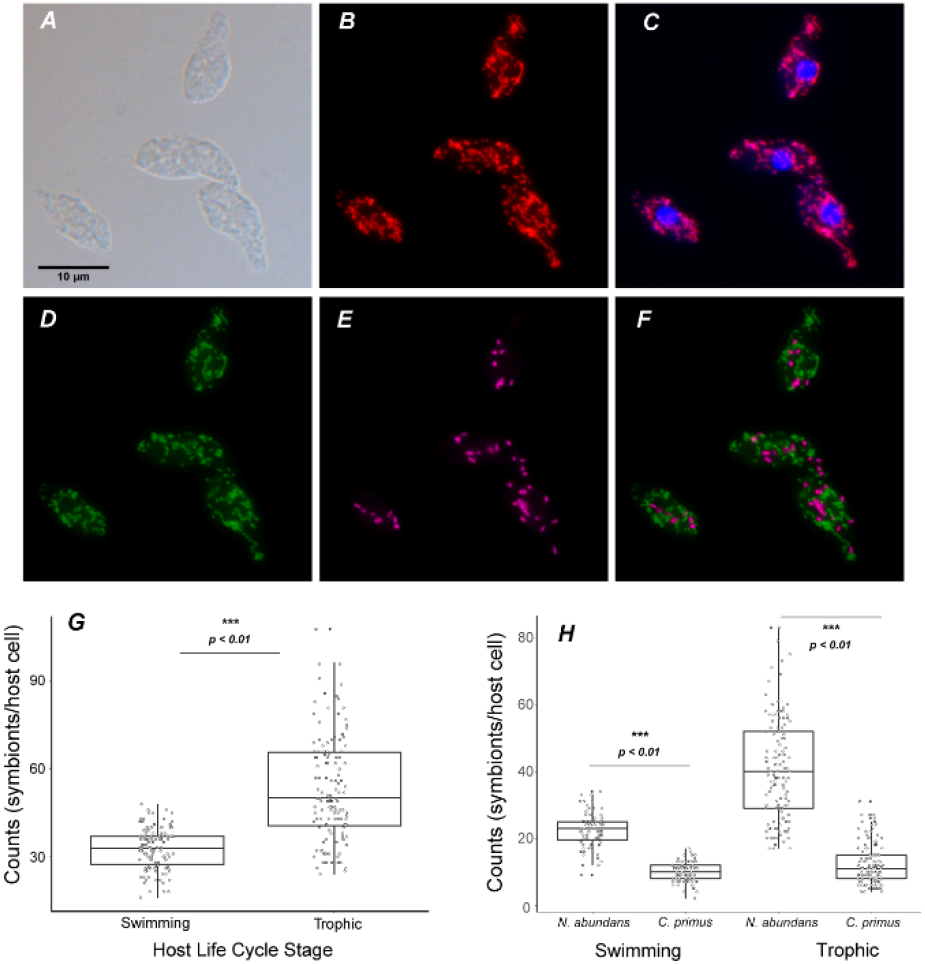
Microscopy and bacterial counts of *D. japonicum* 1604 with *C. primus* and *N. abundans* endosymbionts. Panels A - F show microscopy of diplonemid host cells with endosymbionts. A) DIC; B) FISH-eubacterial probe; C) overlay of DAPI and FISH-eubacterial probe; D) FISH-*C. primus* probe; E) FISH-*N. abundans* probe; F) D and E overlay. G) Total abundance of bacterial endosymbionts was higher in trophic hosts than swimming hosts (p < 0.01). H) *N. abundans* was significantly more abundant than *C. primus* in hosts during both life stages (p < 0.01).

### Diplonemid endosymbionts have small genomes with many uncharacterized proteins

We sequenced the metagenome of each diplonemid, from which we retrieved complete endosymbiont genomes (Star Methods). These genomes were among the smallest recorded for protist endosymbionts. The *Holosporaceae* genomes ranged from 615,988 to 625,897 bp with 29.7 – 30.0% G+C content and 505–550 protein-coding genes. The *S. namystus* genome was composed of two elements, a 605,311 bp chromosome and a 27,632 bp plasmid with 34.9% G+C content and 613 predicted protein-coding genes (Figure 2A). All endosymbiont genomes were gene-dense and with very few pseudogenes or mobile elements (Table S1). The diplonemid endosymbiont genomes shared 223 orthologous genes and *Cytomitobacter* and *N. abundans* genomes shared additional 58 orthologs (Figure 2B). Classifying proteins into Cluster of Orthologous Groups (COGs), we found that the endosymbionts had highly similar relative abundances of most functional categories, with a few exceptions such as motility (Figure 2C). Each genome contained numerous hypothetical proteins, altogether making up 22–50% of the predicted protein coding genes (Table S1), and include many putative secreted proteins which may function in bacterial-host or inter-bacterial interactions.

**Figure 2.**
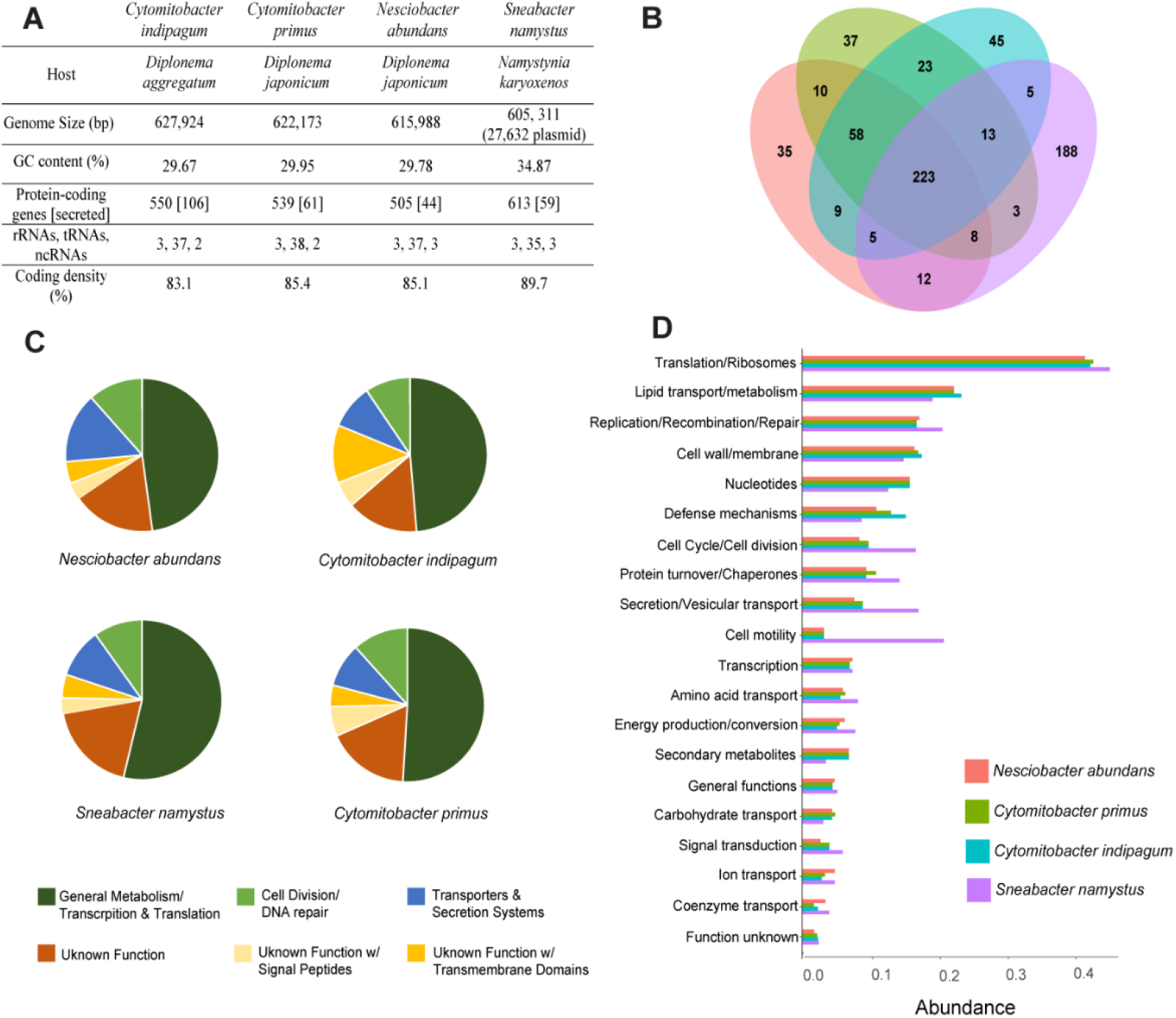
Comparison of diplonemid endosymbiont genomes shows convergent evolution of genome content and function. A) Table of endosymbiont genome content and host species. Secreted proteins were predicted with Phobius and SignalP. See Table S1 for more details. B) *Rickettsiaceae* (*S. namystus*) and *Holosporaceae* (*C. indipagum*, *C. primus, N. abundans*) endosymbionts of diplonemids share 223 orthologous genes and *Holosporaceae* endosymbionts share 58 orthologs. C) Overview of endosymbiont genomes using Pfam annotations and Phobius signal peptide and transmembrane domain predictions. General bacterial metabolism and translation/transcription account for ~50% of the genomes while predicted proteins with unknown functions account for over 25% of the genomes. D) The diplonemid endosymbionts also have similar COG functional category abundances with a few exceptions like motility. COG functional categories were analyzed with web services for metagenomics analysis (WebMGA) and abundances are related to the number of hits to each COG family.

### Symbiont-mediated nutritional provisioning is unlikely

Most highly reduced endosymbionts characterized to date function as nutritional mutualists (Table S2). To determine if the diplonemid endosymbionts were providing their hosts with specific metabolites, we mapped the genomes to the Kyoto Encyclopedia of Genes and Genomes (KEGG) database. All diplonemid endosymbionts had severely reduced energy metabolism with no genes for glycolysis, TCA cycle, or oxidative phosphorylation complexes I–IV (only a partial complex V [ATP synthase] was present) (Figure 3). The *Holosporaceae* diplonemid endosymbionts encoded a partial non-oxidative branch of the pentose phosphate pathway. The retained enzymes of these energy-generating pathways likely serve only for producing biosynthetic intermediates; *N*. *abundans* and *S. namystus* contained a pyruvate dehydrogenase complex that converts pyruvate to acetyl-CoA, and genes for other acetyl-CoA/pyruvate interconversion enzymes were present in all endosymbionts (Figure 3). The most complete metabolic pathways were for the biosynthesis of peptidoglycans, fatty acids, lipids, and iron-sulfur clusters (Figure 3). Other partial pathways in all endosymbionts included myo-inositol biosynthesis and thioredoxin recycling pathways as well as vitamin degradation and salvage pathways. However, there were no complete synthesis pathways for essential metabolites, such as amino acids or vitamins that endosymbionts could provide to their diplonemid hosts.

**Figure 3.**
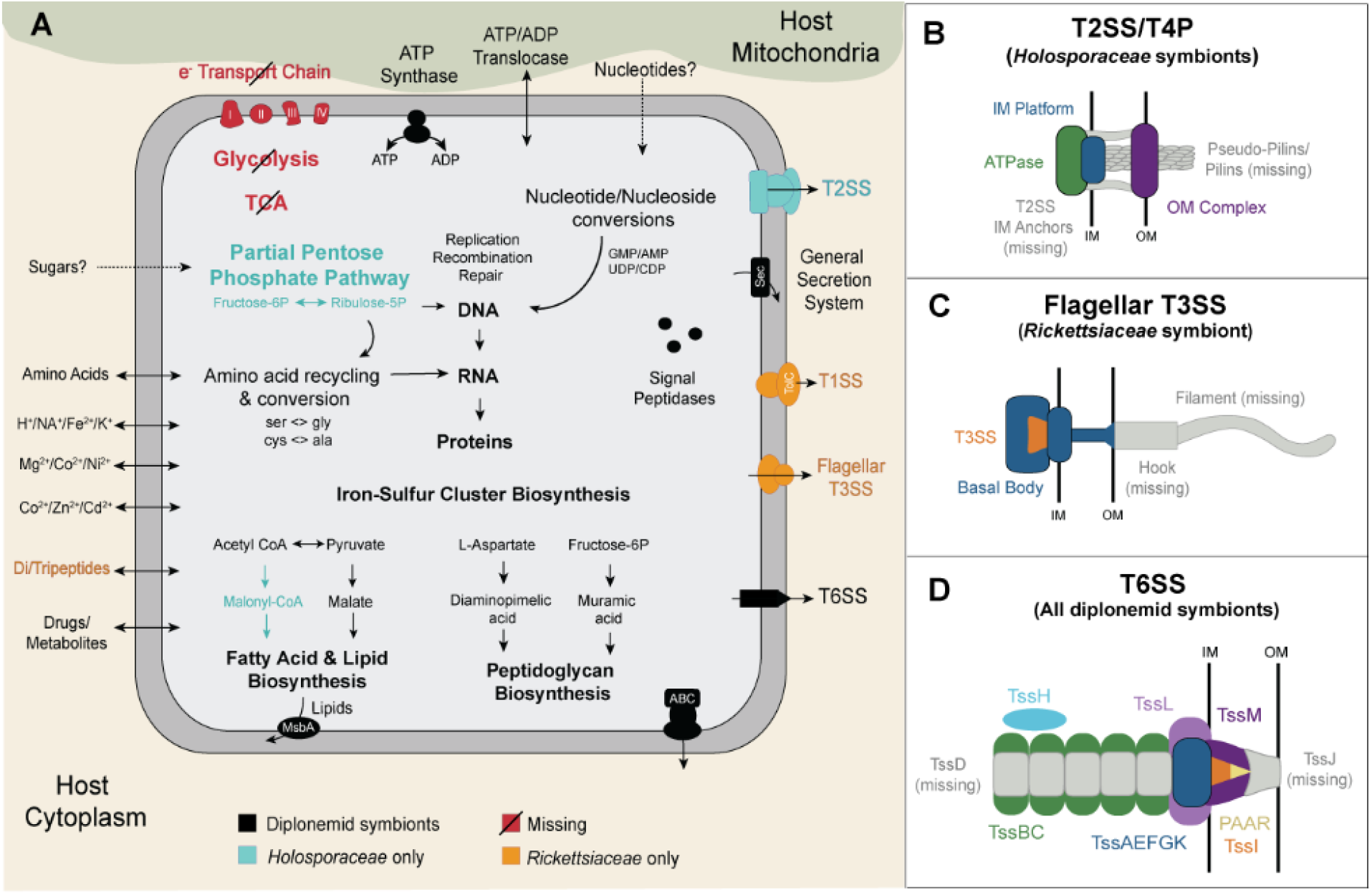
*Rickettsiaceae* and *Holosporaceae* diplonemid endosymbionts have similar reduced metabolic pathways and modified secretion systems. A) Energy pathways are reduced or lost but cellular structure, division and replication pathways have been retained. Nucleotides, amino acids and other metabolites are likely gained from the host via transporters and the endosymbionts encode several copies of the ADP/ATP translocase. Similar toxins and antitoxins are also present and include the T6SS VgrG effector and YwqK immunity proteins. Shared pathways and cellular components are shown in black, missing pathways in red, *Holosporaceae*-specific in blue, and *Rickettsiaceae*-specific in yellow. B – D) Reduced or modified secretion systems of diplonemid endosymbionts. B) *Cytomitobacter* and *Nesciobacter* endosymbionts contain a reduced T2SS/T4P. C) *Sneabacter namystus* encoded a reduced flagellar T3SS with only the basal body present. D) All diplonemid endosymbionts had T6SSs with missing inner tubes and outer membrane components, but an effector, VgrG and immunity protein, TssI, were present. See also Table S3.

Due to their extreme metabolic diminution, the endosymbionts likely depend on the import of many metabolites from their host. Each species encoded between two to four ADP/ATP translocases (*tlc*), which enable the direct import of ATP from the host cytosol. Several other transporters were also present including amino acids and metabolite/drug transporters (Table S3). Considering their lack of pathways for energy production, and their localization near the host mitochondria [11], these endosymbionts may participate in “energy parasitism”, as reported for other members of the *Holosporaceae* and *Rickettsiaceae* [14].

### Numerous secretion systems may mediate intercellular interactions

In contrast to their dramatic metabolic reduction, the diplonemid endosymbiont genomes encoded a large number of genes dedicated to protein secretion. The number of genes with predicted signal peptides (i.e., targeted to membranes or the periplasm) ranged from 44 in *N. abundans* to 106 in *C. indipagum* (Table S1). A reduced general secretion system was present in all diplonemid endosymbionts (Table S3) and a small number of type II secretion system (T2SS) and type IV pili (T4P) genes were found in *Cytomitobacter* and *Nesciobacter* genomes (Figure 3), suggesting a possible T2SS/T4P hybrid secretion system [15]. *Sneabacter namystus* retained a T1SS, and additionally contained a reduced flagellar type III secretion system (T3SS) (Figure 3 and 4). The flagellar basal body was present, but the hook, filament, hook-filament junction, and cap were missing along with several T3SS proteins (Table S3).

**Figure 4.**
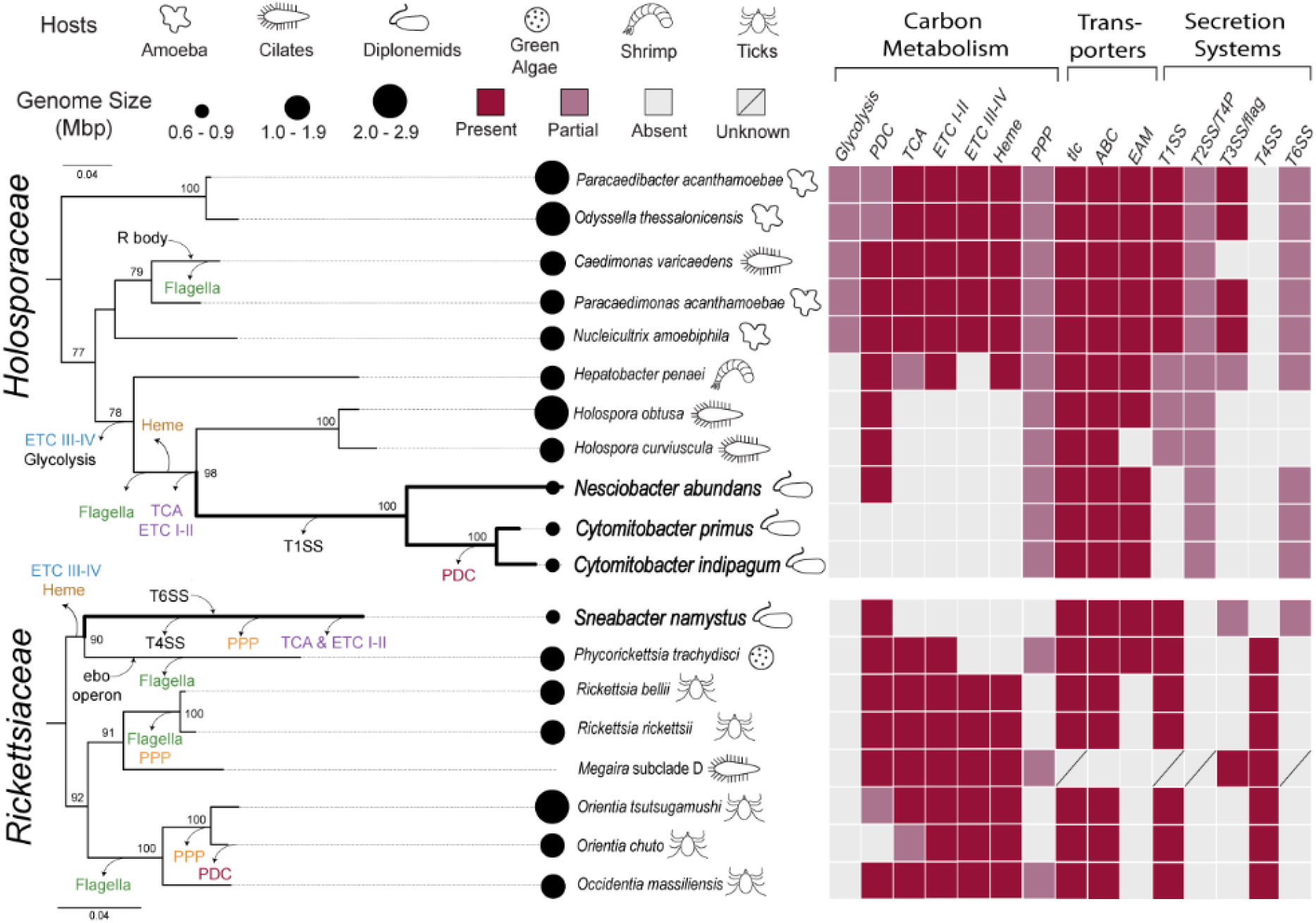
Step-wise reduction and loss of carbon metabolism and secretion systems in both *Holosporaceae* and *Rickettsiaceae* show evolutionary convergence between the two groups of intracellular bacteria. Diplonemid endosymbionts (in bold) from both families of intracellular bacteria have undergone extreme genome reduction with carbohydrate metabolism and secretion system loss (arrows leaving branches). Gain of function has also occurred via horizontal gene transfer such as the T6SS in *S. namystus* and R-body in *C. varicaedens* (arrows pointing to branches). Genome sizes are indicated by black circles and icons depict the host of the endosymbiont. Complete, partial, absent or unknown pathways or secretion systems are indicated by red, pink, light grey and dark grey respectively. Abbreviations include pyruvate dehydrogenase complex (PDC), electron transport chain (ETC), pentose phosphate pathway (PPP), ADP/ATP transporter (tlc), type I secretion system (T1SS), type II secretion system/type 4 pili (T2SS/T4P), type III secretion system/flagella (T3SS/flag), and type 4 and type 6 secretion systems (T4SS, T6SS). Maximum likelihood trees (IQ-TREE) inferred under the TIM3+I+G4 model from full length 16S rRNA and support values represent 100 bootstrap pseudoreplicates (values lower than 65 not shown). See also Figure S4 for additional details on secretion system reduction.

Furthermore, all diplonemid endosymbionts possessed a modified type VI secretion system (T6SS). T6SSs are known for membrane puncturing and toxin delivery in bacterial competition and phagosome evasion [16,17], and hence may play a role in interactions between intracellular endosymbionts, or in the original colonization of the diplonemid hosts. This system appeared to be ancestral to *Cytomitobacter* and *Nesciobacter*, but in *S. namystus*, the highly divergent system was completely encoded on the plasmid. T6SS genes are absent in all other *Rickettsiaceae* genomes (Figure 4), although some T6SS genes were found in a metagenome-assembled *Rickettsiales* genome (Figure S3). Therefore, the T6SS was likely acquired horizontally in *S. namystus* and the acquisition was relatively ancient due to the similar GC content of the plasmid (33.8%) and chromosome (34.9%).

Interestingly, all the diplonemid endosymbionts retained the majority of canonical T6SS components (Figure 3, Table S3), but the inner tube (TssD) and outer membrane complex (TssJ) were missing in all four species, as well as in related *Holosporaceae* (Figure 3, Figure S4), suggesting these bacteria use a specialized, modified T6SS. A TssI (YwqK family) immunity protein was encoded next to the VgrG tip protein in all endosymbionts and the *S. namystus* plasmid contained two VapC toxins encoded next to two antitoxins. Additionally, proteins with leucine-rich repeat (LRR) domains, often associated with eukaryote interactions [18], were present upstream and downstream from T6SS effectors in *Cytomitobacter* and *Nesciobacter*, and other LRRs were found throughout all diplonemid endosymbiont genomes (Table S1).

### Symbiont genomes are relatively stable and retain DNA repair pathways

Genomic erosion can be accelerated by the loss of DNA repair and recombination mechanisms, which can lead to a runaway accumulation of deleterious mutations, i.e., Muller’s Ratchet [19,20]. In the diplonemid endosymbionts, the RecA-dependent RecFOR pathway was present for double strand break repair, but the RecBCD recombination pathway was absent. Other DNA repair mechanisms were also intact: UvrABC and RuvABC in *S. namystus*, and MutSL and DNA polymerase I were identified in all endosymbionts, although several domains were missing in the DNA polymerase I genes. Additionally, the endosymbionts encoded the majority of cell cycle and division genes (*ftsZ*, *ftsW*, *murG*, etc.). The presence of these pathways and the low number of pseudogenes and mobile elements suggest that diplonemid endosymbiont genomes are relatively stable at this point in their evolution despite rapid evolution and GC bias.

### Description of new bacterial species

*Ca*. Nesciobacter abundans gen. nov., sp. nov. belongs to the class *Alphaproteobacteria* [21]; subclass *Caulobacteridae* [22]; order *Rhodospirillales* [23]; and family *Holosporaceae* [24]. The genus name is derived from the Latin word *nescio* meaning ‘unknown’ or ‘I do not know’ and *bacter* referring to bacteria. The species name is derived from the Latin word for abundant and describes the higher abundance of *Ca.* N. abundans compared to the other endosymbiont in the same host cell.

Obligate endosymbionts of *Diplonema japonicum*; reside freely in host cytoplasm short rods [0.9 to 1.2 µm long and 0.5 to 0.7 µm wide]; Gram-negative cell wall organization; flagella absent; granular homogenous electron-dense cytoplasm; no visible inclusions and internal membrane structures. GC content 29.78%. Genus and species assignment based on full-length 16S rRNA gene; deposited under NCBI BioProject PRJNA556273.

## Discussion

Many eukaryotes harbor prokaryotic endosymbionts, but the study of these – and particularly those with the most severely reduced genomes – has been historically biased towards those found in animal hosts [4,25]. Yet, the most organelle-like endosymbiont is known from a protist host [26,27]. The functional nature of these animal-bacterial symbioses is also biased: of the approximately 210 endosymbionts with sequenced genomes under 1 Mb, 87% are engaged in nutritional mutualisms (Table S2). Therefore, a great deal of the theoretical basis for our understanding of the impacts of endosymbiosis are based on these systems, but the scope of microbial symbioses at the lower limits of cellular and genomic complexity are likely much more diverse and more ambiguous than has been found thus far.

For example, the *Kinetoplastibacterium* spp. endosymbionts (742–833 kb) of trypanosomatids aid in the synthesis of heme and amino acids [28], retaining many metabolic genes. In contrast, the recently discovered endosymbiont of a *Euplotes* ciliate, *Pinguicoccus supinus* (at 163 kb, the smallest protist endosymbiont genome to date) lacks most metabolic pathways, but interacts closely with host lipid droplets, for unknown reasons [29]. *Fokinia solitaria* (837 kb) is another mysterious protist endosymbiont [30], which bears many similarities to the diplonemid endosymbionts; these include the lack of many central metabolic pathways, the presence of ADP/ATP translocases, and a diverse arsenal of protein secretion systems [30]. These commonalities indicate that particular intracellular lifestyles can be converged upon from different evolutionary starting points. Indeed, the diplonemid endosymbionts have converged on very similar genome sizes and contents starting from two distantly related bacterial lineages.

Comparing the genomes of diplonemid *Holosporaceae* and *Rickettsiaceae* to their close relatives, we find overarching patterns of convergence in the step-wise reduction and loss of carbon metabolism and secretion systems (Figure 4). The absence of glycolysis is well known in *Rickettsiaceae* where the endosymbionts import metabolites from their host [31], but loss of glycolysis was only recently discovered in certain *Holosporaceae* species [32]. Although most *Holosporaceae* genomes encoded reduced glycolytic pathways, a clade containing the diplonemid endosymbionts, *Holospora* and *Hepatobacter* had completely lost glycolysis along with the respiratory chain complexes III and IV (Figure 4). Oxidative phosphorylation was even further reduced in both *Holosporaceae* and *Rickettsiaceae* diplonemid endosymbionts as well as in *Holospora*, where the loss of the TCA cycle coincided with the absence of NADH dehydrogenase (complex I) and succinate dehydrogenase (complex II). Despite the absence of the respiratory chain, ATP synthase was retained in all diplonemid endosymbionts and this pattern has been observed in other highly reduced endosymbionts of various hosts [1,33], where it may be used to hydrolyze ATP to generate a proton gradient [34,35]. This extreme reduction of energy and carbon metabolism in diplonemid endosymbionts compared to their *Holosporaceae* and *Rickettsiaceae* relatives increased the endosymbionts reliance on diplonemid metabolites and likely led to additional metabolic and genomic reduction.

Lineage-specific flagellar reduction and loss is also common to both families: flagella have been lost at least three times independently in *Rickettsiaceae* and at least twice in *Holosporaceae*. In spite of the flagellar reduction in both groups of diplonemid endosymbionts (Figure 4), the flagellar basal body and T3SS were found in *S. namystus*. This has also been observed in the *Buchnera* endosymbionts of aphids [36–38], where flagellar basal bodies cover the surface [37] and have likely been repurposed for secretion of unknown effectors instead of flagellin [39,40], which is also the most plausible function in *S. namystus*. Therefore, the reduced flagellar T3SS is an example of potentially repurposed cellular machinery in intracellular bacteria and of evolutionary convergence in unrelated endosymbionts of protists and animals.

Indeed, a large portion of the reduced diplonemid endosymbiont genomes are dedicated to protein secretion, including the presence of an extensive arsenal of effectors and putative toxins. While prominent proportionally, the number of secretion systems were diminished compared to relatives with larger genomes, and tended to have smaller or modified gene complements (Figure 4, Figure S4). For example, two major T6SS components, the inner tube and outer membrane complex, were absent in the diplonemid endosymbionts along with many other *Holosporaceae* endosymbionts, but also in *S. namystus* (Figure 3 and 4). These components are required for the extension and membrane-puncturing capability of the T6SS in bacterial competition and phagosome evasion [16], but those T6SS functions may not be necessary for intracellular endosymbionts that live outside host vesicles. Several leucine-rich repeat (LRR) domains were also encoded next to T6SS proteins and LRRs are known to be involved in interactions with eukaryotic proteins [41,42]. Additionally, the acquisition of plasmid-based T6SS from an unknown source in *S. namystus* (Figure S3) makes the *Rickettsiaceae* endosymbiont more similar in general capabilities to *Holosporaceae* endosymbionts, and appears important to the establishment of symbiosis in diplonemids (Figure 4). This and other horizontal gene transfers present in *Holosporaceae* [43] and *Rickettsiaceae* [44] protist endosymbionts likely contributed to their intracellular specialization to specific hosts (Figure 4).

The functions of secreted effectors and putative antimicrobial toxins are largely unknown in the *Holosporaceae* and the *Rickettsiaceae*. The diplonemid endosymbionts have been found to associate with host mitochondria [11] and the bacteria might use effectors to interact with the mitochondria for ATP acquisition, or with cytoskeletal proteins to mediate their spatial positioning within the host. The retention of effector and immunity protein pairs along with secretion systems hints at the critical role of cellular interactions initiated by the endosymbionts, but the target of these manipulations remains unclear. Secreted effectors are probably used against the host to establish stable colonization, but some may also be targeting the mitochondria, or even other intracellular bacteria. They could be used to signal conspecifics, compete with other co-occurring endosymbiont species, or repel foreign invaders [45]. In the latter case, this may be beneficial to the diplonemid host, as it could protect it against infection by bacterial parasites. Examples of bacterial defensive symbiosis have been found in *Acanthamoeba* protists, where endosymbionts can prevent the initial entry of the pathogen or interfere with its life cycle within the host [5,46,47]. If the diplonemid endosymbionts are beneficial, other functions they may provide include osmotic stress resistance [48], heat tolerance [49] or even nutritional supplementation that is not yet obvious, given the large number of uncharacterized proteins encoded in their genomes (Figure 2). Yet, the endosymbionts may also be parasites that are no longer able to spread through horizontal transmission, leading to further reduction of their genomes.

The co-occurrence of two different *Holosporaceae* endosymbionts in *Diplonema japonicum* (Figure 1) raises further questions about diplonemid endosymbiont function, conflict, and transmission. Most *Diplonema* species lack endosymbionts [11,12] and even with the limited sampling now available, *C. primus* and *N. abundans* are not sister species, altogether arguing against the conclusion that they speciated within *D. japonicum*. The secretion of T6SS effectors would generally exclude colonization by new endosymbionts (if they are experimentally verified to be toxin-antitoxin systems as seems likely), but the presence of similar immunity proteins in all diplonemid endosymbionts may relax such restrictions, allowing ancestors of either *C. primus* or *N. abundans* to inhabit a host already infected with the other. In animal systems, such co-occurrence has been observed to lead to partitioning of essential functions [1,3], but this is most obviously applicable to nutritional supplementation and there is no evidence for the partitioning of any pathway in *C. primus* and *N. abundans*. The co-occurrence of a nutritional symbiont with a defensive symbiont has also been found in animal systems [50], but if the diplonemid symbionts are defensive, then this would be the first case of two defensive symbionts in one host.

All the diplonemid endosymbionts, along with other reduced *Rickettsiaceae* and *Holosporaceae* symbionts, are susceptible to the evolutionary consequences of genome reduction and likely experience population bottlenecks due to factors like host cell division and starvation, or endosymbiont life stages. The decreased endosymbiont load in starving diplonemid hosts (Figure 1) suggests that small endosymbiont populations occur in nature and a sharp reduction in population size can lead to the inevitable fixation of deleterious mutations [19,20]. The retained recombination and repair mechanisms in *Rickettsiaceae* and *Holosporaceae* endosymbionts may correct deleterious mutations, thus slowing Muller’s ratchet [19], but why the highly reduced diplonemids have retained the mechanisms while other highly reduced endosymbionts have lost them is yet to be understood [51]. Additional studies on endosymbiont populations in both protist and animal hosts will provide insight into the mechanisms of stable symbiosis and distinguish context-specific characteristics from more general principles of endosymbiosis.

Overall, the genome reduction of *Rickettsiaceae* and *Holosporaceae* diplonemid endosymbionts has converged on very similar genome content and metabolic potential, and suggests that extremely streamlined endosymbionts can be common associates of many single-celled eukaryotes. Several of these characteristics are common to other *Rickettsiaceae* and *Holosporaceae* endosymbionts, but the diplonemid endosymbionts have all evolved to greater extremes in parallel. The commonalities between the diplonemid endosymbionts and unrelated endosymbionts of animals indicate that particular cellular machinery such as the retention of modified secretion systems (e.g., flagellar basal bodies) and ATP synthase without oxidative phosphorylation is advantageous in a large range of hosts. Although these modified cellular features have been observed before, they have never been found in a single host-symbiont system. Therefore, this study provides a model of convergent evolution in endosymbionts and reveals the invisible interactions between bacteria and one of the most abundant marine protists.

## Acknowledgments

We thank Dr. Vittorio Boscaro for his helpful advice and knowledge on endosymbionts of protists. We also thank Dr. Sergio Muñoz-Gómez for the multi-protein alignment. Research was supported by the Moore Foundation. E.E.G. was supported by the UBC International Fellowship, F.H. was supported by the EMBO Long-Term Fellowship and W.K.K was funded by the Killam Postdoctoral Research Fellowship.

## Author Contributions

Genome Sequencing, E.E.G., F.H., D.T., G.P. and A.H.; Genome Analysis E.E.G, F.H. and W.K.K.; Culture Establishment and FISH Experiments, D.T. and G.P.; Writing E.E.G., F.H., W.K.K. and P.K. All authors contributed to the revisions of the manuscript. Funding Acquisition and Supervision, J.L. and P.K.

## Materials and Methods

### Cultures

Five different species/strains of diplonemids containing endosymbionts were analyzed: *Diplonema japonicum* YPF1603 and YPF1604, *D. aggregatum* YPF1605 and YPF1606, and *Namystynia karyoxenos* YPF 1621. *Diplonema* spp. were grown axenically in seawater-based Hemi medium at 15 °C according to Tashyreva et al., 2018. *Namystynia karyoxenos* was grown at 21°C in 12 hour light and dark cycles. For DNA extraction, diplonemid cultures were grown to a maximum concentration of 1×10^5^ cells/ml to 7×10^5^ cells/ml.

### DNA extraction, genome sequencing and assembly

The genomes of four diplonemid endosymbionts sequenced included *Candidatus* Cytomitobacter primus, *Ca*. Cytomitobacter indipagum, *Ca*. Nesciobacter abundans (*Holosporaceae*), and *Ca.* Sneabacter namystus (*Rickettsiaceae*) that will be referred to without the *Candidatus* prefix. A QIAGEN Power Biofilm kit was used for DNA extractions and the quality and quantity of each sample was recorded by Nandrop and Quibit readings. DNA library preparations were performed with the Nextera XT and TruSeq library kits (*N*. *abundans* and *C*. *primus*) and sequenced using Illumina MiSeq (*N*. *abundans* and *C. primus*) and Illumina Hiseq 2500 (all other species). The type of reads used for Hiseq was 2 x 125 bp and average coverage was 69x (*S*. *namystus*), 164x (*C. primus*), 310x (*N. abundans*) and 3423x (*C. indipagum*).

Each genome was assembled in SPAdes v3.11.1 [52] and host and containment contigs were removed from endosymbiont contigs in Blobtools v1.0.1 [53] using GC content and coverage thresholds. In addition, Oxford Nanopore MinIon 1D ligation library was sequenced for *S*. *namystus* and *C. indipagum* and these genomes were closed using Unicycler v0.4.7 [54]. PROKKA v1.12 [55] and RAST server [rast.nmpdr.org] were used for functional annotation. Orthologous gene comparison was conducted with EggNOG v4.5.1 [56] and protein family annotation was conducted with the Pfam v31 database using a Hidden Markov Model (HMM) search.

Maximum likelihood trees of bacterial 16S rRNA and other bacterial genes were made in IQ-TREE v1.5.4 and metabolic pathways were constructed in Pathway Tools [57] and the KEGG Automated Annotation Server [genome.jp/kegg/kaas]. Secretion systems were identified with BLAST [58] and TXSScan [59], and type VI secretion systems were further analyzed using the SecReT6 database [60]. Signal peptides and transmembrane domains were found with the Phobius webserver [phobius.sbc.su.se] [61]. Pseudogenes were estimated using Pseudofinder [github.com/filip-husnik/pseudo-finder] [62]. Genome trait comparisons were conducted using BLAST and all *Rickettsiales* and *Rhodospirillales* genomes used in the comparisons were publicly available and downloaded from the National Center for Biotechnology Information (NCBI) database.

Both *Cytomitobacter* species had been previously described, and included *C. primus* found in *D. japonicum* YPF1604 and *C. indipagum* found in *D. aggregatum* YPF1606 [11]. Endosymbiont genomes from clonal *D. japonicum* YPF1603 and *D. aggregatum* YPF1605 strains were also assembled but the genomes had nearly identical sequences to *C. primus*, *N*. *abundans* and *C. indipagum* (Supplementary Figure 1), and were not analyzed further.

### Fluorescence in situ hybridization (FISH) and DNA staining

The FISH probes for distinguishing bacterial endosymbionts were designed on the basis of full-length 16S rRNA sequences: *N. abundans-*specific HHC117 (5’-CCCTCCATATGGCAGATTCCC-3’) and HLC36 (5’-CATGTGTTAAGCGCGCCGC-3’) specific to *C. primus*, were 5’-labelled with FITC and Cy5 fluorescent dyes, respectively. In addition, Eub338 probe targeting most groups of bacteria (5’-GCTGCCTCCCGTAGGAGT-3’) 5’-labelled with Cy3, was used for total bacteria counts. The probes were designed solely for distinguishing the endosymbionts in *D. japonicum*, and were not tested against any 16S rRNA database. The specificity and efficiency of HHC117 and HLC56 probes were confirmed *in silico* using mathfish.cee.wisc.edu tools and experimentally tested by hybridization in buffers containing 20%, 25%, 30% and 35% (v/v) formamide. Hybridization efficiency was improved by the addition of unlabeled helper oligonucleotides targeting the 16S rRNA regions adjacent to both 3’ and 5’ ends of HHC117 and HLC36 probes.

Both trophic and swimming cells were cultured in triplicates. Cell pellets were fixed with 4% paraformaldehyde in seawater for 30 min, rinsed with dH2O and air-dried on poly-L-lysine-coated glass slides. Adhered cells were dehydrated with 50%, 80% and 96% ethanol solutions for 3 min each. The slides were incubated simultaneously with 250 nM Eub338, HHC117 and HLC36 probes in hybridization buffer (900 mM NaCl, 20 mM Tris/HCl, 0.01% SDS) containing 20% (v/v) formamide at 46°C for 2 hours. The probes were removed by incubation in washing buffer (225 mM NaCl, 20 mM Tris/HCl, 0.01% SDS) on a shaker at 48°C for 30 min. FISH-labelled samples were air-dried and mounted in ProLong Gold antifade reagent (Life Technologies) containing 4’,6-diamidino-2-phenylindole (DAPI).

FISH-labelled slides were viewed with the AxioPlan 2 fluorescence microscope (Zeiss, Germany) equipped with differential interference contrast (DIC) and Chroma F31-01 (FITC), F31-002 (Cy3) and F41-008 (Cy5) filters. Images were taken with DP72 microscope digital camera at 1600×1200-pixel resolution using CellSens software v. 1.11 (Olympus) and processed with GIMP v. 2.8.14, Irfan view v. 4.41 and Image J v. 1.51 software. In each replicate, the number of *N. abundans* and *C. primus* endosymbionts were counted in 35 to 50 cells. The statistical analyses were conducted in RStudio v3.4.2 using simple t-test analyses in the R Stats Package. The fluorescence images were produced by overlaying 2 to 6 images focused on different planes of cells.

