## Supplemental Information for "Highly reduced genomes of protist endosymbionts show evolutionary convergence"

**Supplemental Data Items**

Table S1: Number of predicted diplonemid endosymbiont genome features. Related to Figure 2.

| | *C. primus* | *C. indipagum* | *N. abundans* | *S. namystus* |
|---|---|---|---|---|
| LRR_8 | 3 | 22 | 0 | 3 |
| LRR_4 | 4 | 4 | 2 | 2 |
| Signal peptides | 60 | 106 | 44 | 63 |
| Hypothetical proteins | 169 | 276 | 116 | 155 |
| Phage genes | 5 | 5 | 5 | 8 |
| Pseudogenes | 7 | 16 | 8 | 9 |

Table S2: GenBank endosymbiont genomes smaller than 1 Mb. Related to Figure 2.

| Endosymbionts and host group | | Symbiosis Function | |
|---|---|---|---|
| Total Genomes; Total Genera | 210; 43 | Nutritional | 178 |
| Animal host | 184 | Defensive | 3 |
| Protist host | 10 | Parasitic | 13 |
| Unknown host | 8 | Unknown | 16 |

Table S3: List of secretion system and transporter genes present in endosymbionts. Related to Figure 3.

| | *C. primus* | *C. indipagum* | *N. abundans* | *S. namystus* |
|---|---|---|---|---|
| IM Translocase | *yidC*, *yajC*, *ftsY* | *yidC*, *yajC*, *ftsY* | *yidC*, *yajC*, *ftsY* | *yidC*, *yajC*, *ftsY* |
| General Secretion | *secABDEFY*, *ffh* | *secABDFY*, *ffh* | *secABDEFY*, *ffh* | *secABDFY*, *ffh* |
| T1SS | None | None | None | *hlyB*, *hlyD*, *tolC* |
| T2SS/T4P | *gspF, gspD, pilB, pilQ* | *gspF, gspD, pilM, pilQ, pilB* | *gspF, gspD, pilB, pilQ* | None |
| T3SS/Flagella | None | None | None | *flbB, flgABCFGHI, flhAB, motAB fliFGILMNPQR,* |
| T6SS | *tssABCEFGIKLM tssI/vgrG, cts2N, iglG*/PAAR | *tssABCEFGHIKLM, tssI/vgrG*, *cts2N, iglG*/PAAR | *tssBCEFGIKLM*, *tssI/vgrG*, *cts2N, iglG*/PAAR | *tssABCEFGHKLM, tssI/vgrG*, *iglG*/PAAR, |
| Tol-Pal | *yidD* | *tolA* | *ybgF* | *tolC*, *hlyB*, *hlyD* |
| ATP/NTT transporters | *tlc1* (3) | *tlc1* (2) | *tlc1* (3) | *tlc1, tlc2, tlc4, tlc5* |
| Amino Acid transporters | *potE*, *cadB*, *eamA/rhaT* | *eamA/rhaT* | *eamA/rhaT* | *eamA/rhaT* |
| Other transporters | *proP, mgtE, kefAB,* ABC transporter, *znuC*, RND multidrug efflux transporter | *proP, kefAB, mgtE, znuC*, RND multidrug efflux transporter, ABC transporter | *proP, mgtE*, *kefC*, ABC transporter, *bcr*/*cflA*, *adiC* | *prop, mgtE*, *kefB, nhaA*, *ntrY, znuB,* ABC transporter |

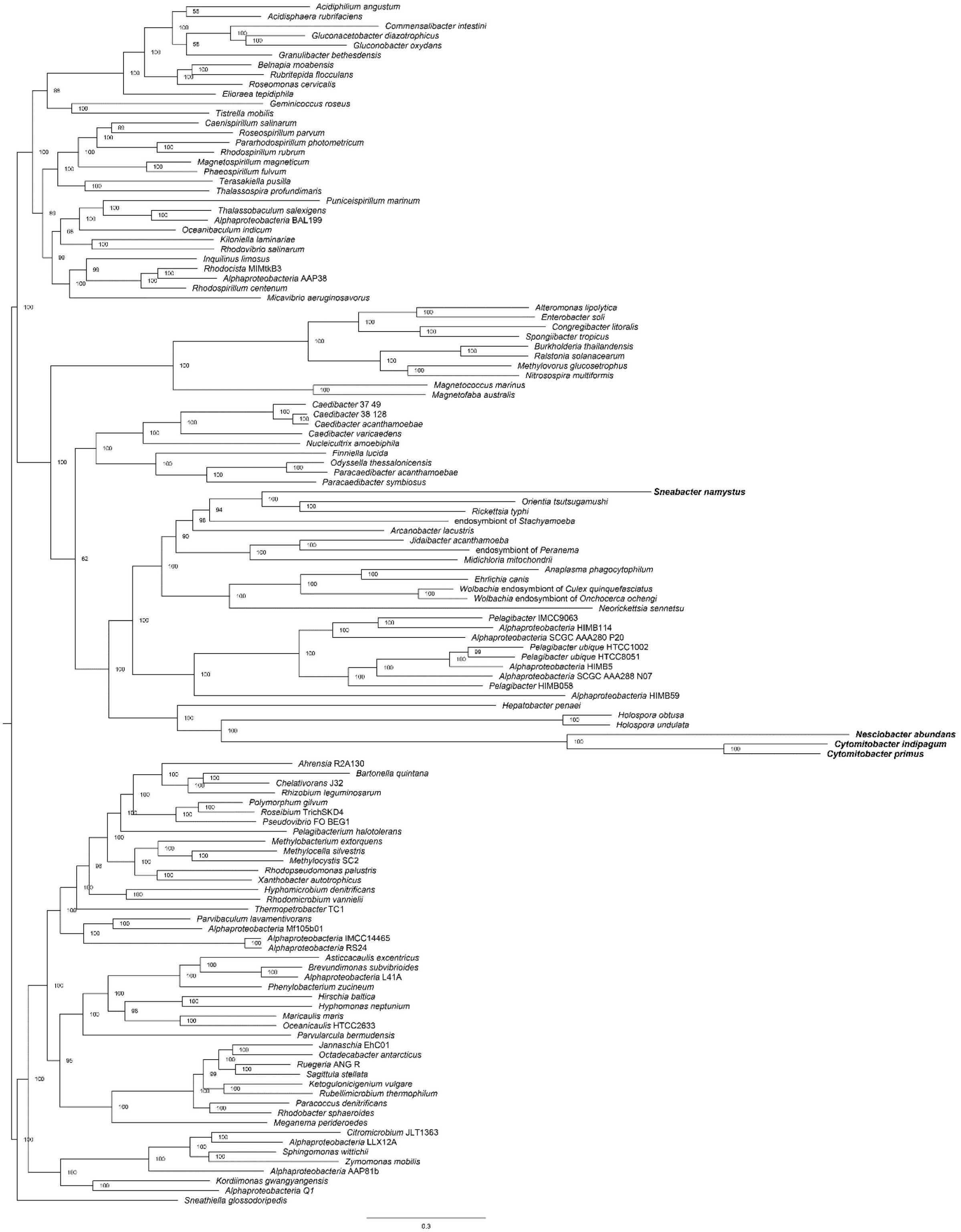


Figure S1. Maximum likelihood trees (IQ-TREE) inferred under the best fitting model (LG+F+R10) from a multi-protein alignment of 200 proteins used in Muñoz-Gómez et al., 2019. Related to Figure 4. Diplonemid symbionts (bold) are well supported within *Rickettsiaceae* and *Holosporaceae*, but the two families group together due to the presence of rapidly evolving sites [1].

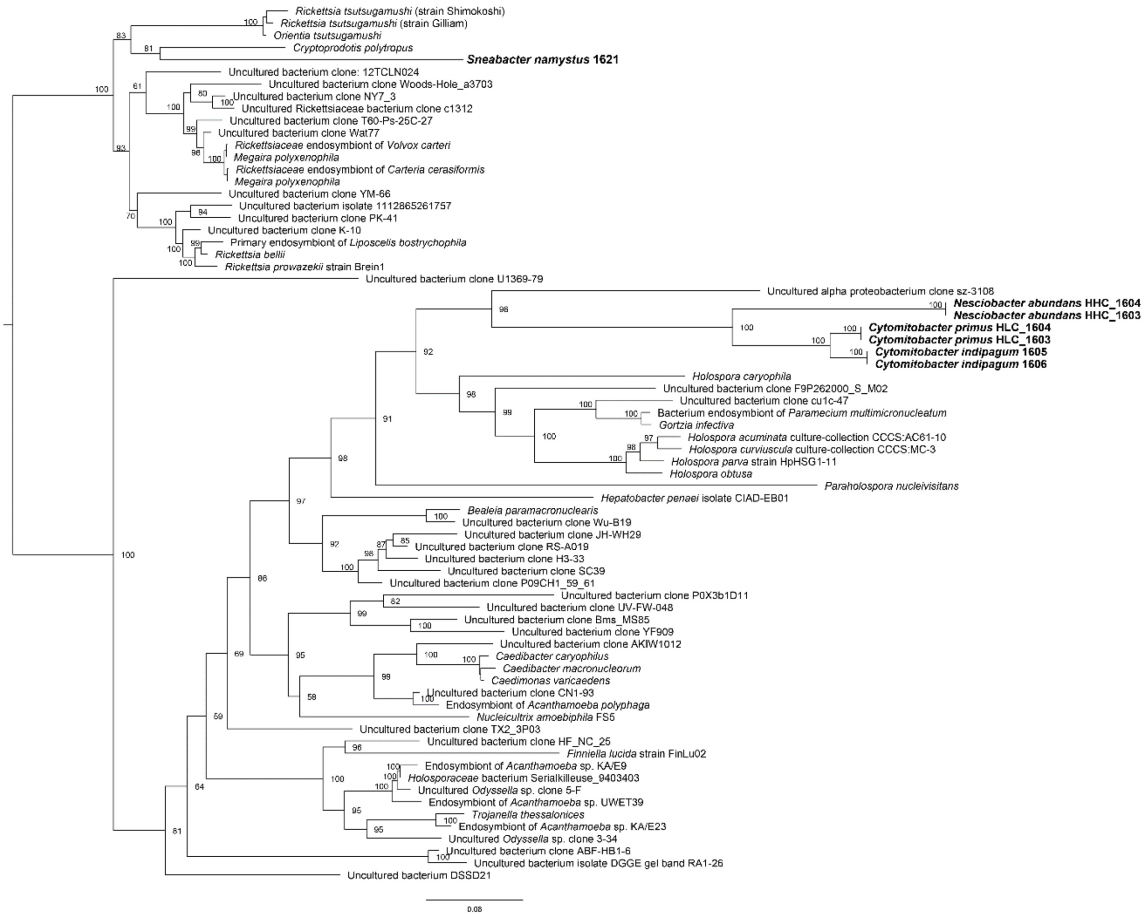


Figure S2. Maximum likelihood tree inferred under the TIM2+I+G4 model from the 16S rRNA gene sequences of diplonemid endosymbionts and related bacterial species. Related to Figure 4.

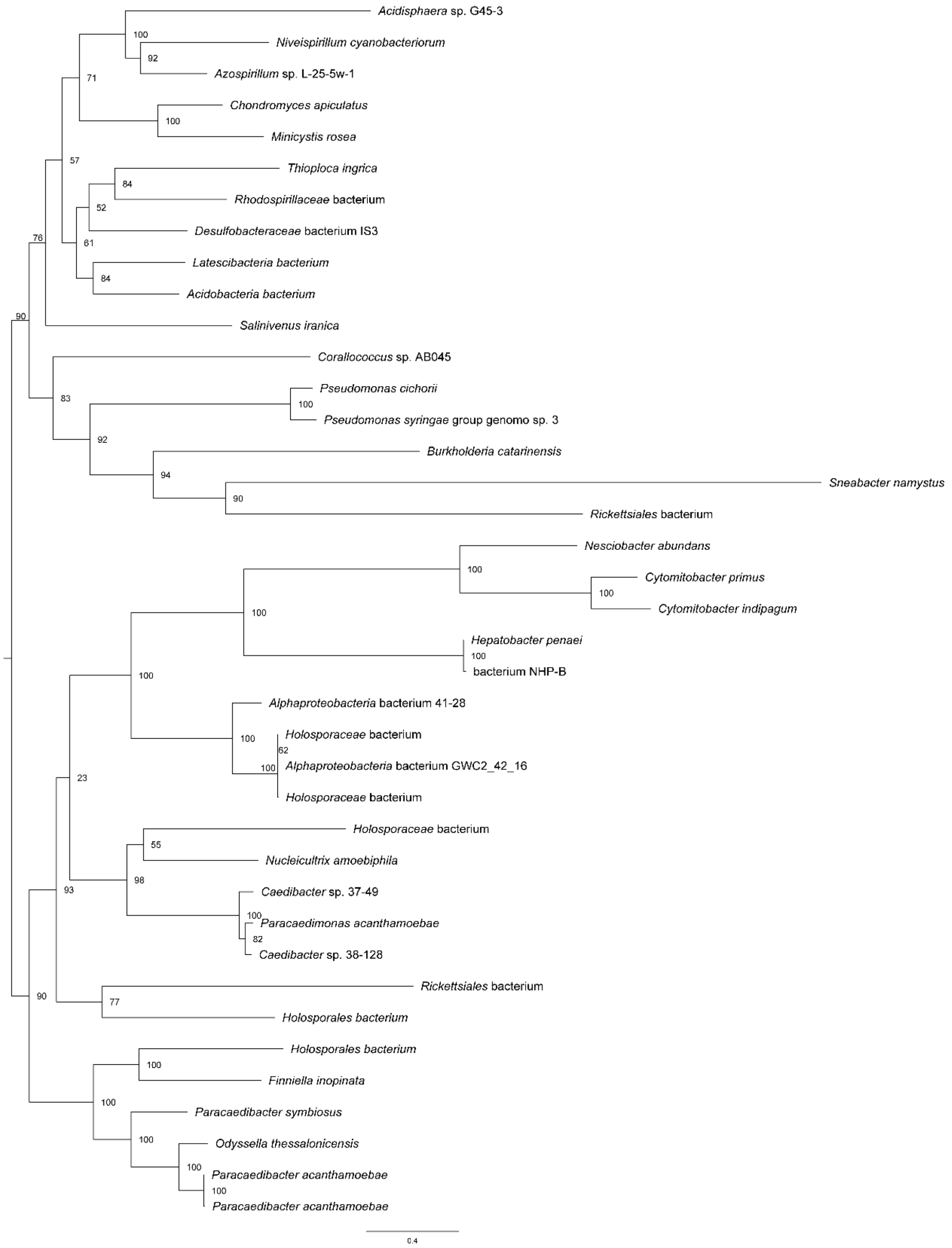


Figure S3. Maximum likelihood tree inferred under the LG+I+G4 model from the type VI secretion system TssF proteins of diplonemid endosymbionts and other bacterial species. Related to Figure 3 and Figure 4.

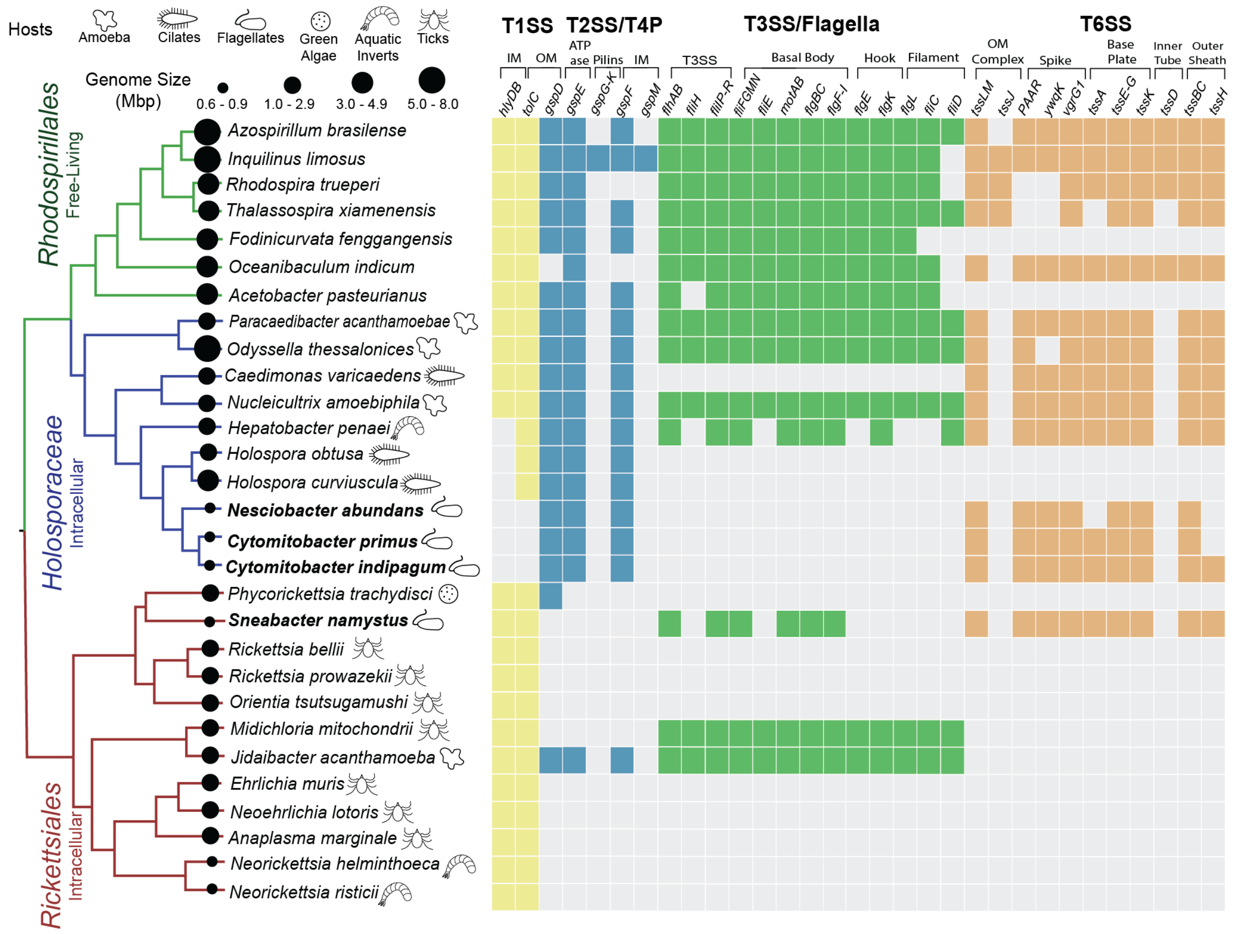


Figure S4. Modified and reduced secretion systems are found throughout *Alphaproteobacteria*. Related to Figure 4. Presence and absence of genes for Type I, II/IVP, III/Flagella and VI Secretion Systems in *Rickettsiales* (red), *Holosporaceae* (blue, family in *Rhodospirillales*), and other *Rhodospirillales* (green) taxa are depicted by colored or gray boxes. All *Rickettsiales* taxa here are endosymbionts of protists and animals while *Rhodospirillales* taxa range from endosymbionts to free-living bacteria. Genome sizes are indicated by black circles and host icons correlate with endosymbionts.
